# Metatranscriptome analysis of the vaginal microbiota reveals potential mechanisms for protection against metronidazole in bacterial vaginosis

**DOI:** 10.1101/248302

**Authors:** Zhi-Luo Deng, Cornelia Gottschick, Sabin Bhuju, Clarissa Masur, Christoph Abels, Irene Wagner-Döbler

## Abstract

Bacterial vaginosis (BV) is a prevalent multifactorial disease of women in their reproductive years characterized by a shift from the *Lactobacillus* spp. dominated microbial community towards a taxonomically diverse anaerobic community. For unknown reasons, some women do not respond to therapy. In our recent clinical study, out of 37 women diagnosed with BV, 31 were successfully treated with metronidazole, while 6 still had BV after treatment. To discover possible reasons for the lack of response in those patients, we performed a metatranscriptome analysis of their vaginal microbiota, comparing them to patients who responded. Seven out of 8 Cas genes of *Gardnerella vaginalis* were highly up-regulated in non-responding patients. Cas genes, in addition to protecting against phages, might be involved in DNA repair thus mitigating the bactericidal effect of DNA damaging agents like metronidazole. In the second part of our study, we analyzed the vaginal metatranscriptomes of four patients over three months and showed high *in vivo* expression of genes for pore-forming toxins in *L. iners* and of genes encoding enzymes for the production of hydrogen peroxide and D-lactate in *L. crispatus*.

## Importance

Bacterial vaginosis is a serious issue for women in their reproductive years. Although it can usually be cured by antibiotics, the recurrence rate is very high, and some women do not respond to antibiotic therapy. The reasons for that are not known. Therefore we undertook a study to detect the activity of the complete microbiota in the vaginal fluid of women that responded to antibiotic therapy and compared it to the activity of the microbiota in women that did not respond. We found that one of the most important pathogens in bacterial vaginosis, *Gardnerella vaginalis*, has activated genes that can repair the DNA damage caused by the antibiotic in those women that do not respond to therapy. Suppressing these genes might be a possibility to improve the antibiotic therapy of bacterial vaginosis.

## Introduction

The healthy vaginal microbiome is characterized by low pH and low diversity and can be categorized into community state types (CSTs) that are dominated by different *Lactobacillus* spp. such as *L. crispatus*, *L. iners*, *L. gasseri* and less frequently *L. jensenii* or a more diverse community (1). Bacterial vaginosis (BV) is a frequent multifactorial disease of women in their reproductive years that is characterized by a shift of this *Lactobacillus* spp. dominated bacterial community to a community of various mostly anaerobic bacteria (2). BV is associated with a higher risk of preterm birth and of acquiring sexually transmitted infections such as HIV (3). The most common bacteria found in BV, identified by 16S rRNA gene sequencing, are *Gardnerella*, *Atopobium*, *Prevotella*, *Bacteroides*, *Peptostreptococcus*, *Mobiluncus*, *Sneathia*, *Leptotrichia*, *Mycoplasma* and BV associated bacterium 1 (BVAB1) to BVAB3 of the order Clostridiales. Recently, three CSTs dominated by *Gardnerella vaginalis*, Lachnospiraceae and *Sneathia sanguinegens*, respectively, have been described (4). In our recent clinical study *S. amnii* was identified as the best biomarker for BV (5).

The most important pathogen in BV is *Gardnerella vaginalis* (6). It is currently the only described species in the genus *Gardnerella*, but genome comparisons suggest that it can be separated into four genetically isolated subspecies (7). While they cannot be resolved by 16S rRNA gene sequencing, the universal target from the chaperonin-60 gene separates the species into the same four subgroups (group A, clade 4; subgroup B, clade 2; subgroup C, clade 1; subgroup D, clade 3) (6,8,9). All four subgroups of *G. vaginalis* can be detected in the vaginal microbiota of healthy women throughout the menstrual cycle (10). Subgroup A and C define distinct CSTs in health (11). Isolates from the four subgroups of *G. vaginalis* differ in their virulence as well as in their resistance against metronidazole. The sialidase activity of *G. vaginalis* is an important virulence factor and it was detected in all isolates from subgroup B and few isolates of subgroup C but not in subgroups A and D isolates (12). The presence of sialidase activity is used for diagnosis of BV in a commercial kit (13). Resistance against metronidazole was found in subgroups A and D isolates, while those from subgroups B and C were highly susceptible (14).

Metronidazole is a widely applied chemotherapeutic agent used to treat infectious diseases caused by anaerobic bacteria, and it is the first-line antibiotic for treating BV (15,16). Metronidazole is a prodrug which requires enzymatic reduction within the cell, which occurs under anaerobic conditions only, to transform it into an active form (17). Activated metronidazole acts by covalently binding to DNA, disrupting its helical structure and causing single and double strand breaks that lead to DNA degradation and death of the pathogens (17). Resistance can therefore be mediated by lack of activation of the prodrug, or by repair of DNA damage, and has been studied in various pathogens. In *Helicobacter pylori* and *Campylobacter spp*. ferredoxin, ferredoxin/ferredoxin-NADP reductase (FNR) and nitroreductase contribute to metronidazole resistance (17). In *Bacteroides fragilis*, genes responsible for DNA repair like recA and recA-mediated autopeptidase (Rma) and a gene named nitroimidazole resistance gene (*nim*) encoding a nitroimidazole reductase were shown to confer resistance against metronidazole (18,19). Failure of BV treatment by metronidazole is relatively rare (5,20). It is unclear if it is caused by resistance of the BV pathogens to metronidazole, and which mechanisms are acting *in vivo*. A recent study has demonstrated that failure of treatment of BV with metronidazole is not associated with higher loads of *G. vaginalis* and *A. vaginae* (21). Isolates from *G. vaginalis* subgroups A and D are intrinsically resistant against metronidazole, but the underlying mechanism is unknown (14).

Until now, the majority of studies regarding the vaginal microbiota have focused on 16S rRNA gene sequencing, answering only questions on the taxonomic composition of bacterial communities but not on their functions (2). A metatranscriptome analysis comparing vaginal swabs from two women with BV with two healthy subjects showed that *L. iners* upregulates transcription of the cholesterol-dependent cytolysin (CDC) and of genes belonging to the clustered regularly interspaced short palindromic repeats (CRISPR) system in BV (22). No study has investigated the activity shifts of the vaginal microbiota during antibiotic treatment of BV.

We had previously analyzed the vaginal microbiota in the context of a clinical trial using 16S rRNA gene sequencing (5). Of 37 patients diagnosed with BV and included in this study, 31 were initially cured by a single oral dose of metronidazole. Six patients did not respond, i.e. they were still diagnosed with BV according to Nugent score after antibiotic therapy. Here we asked if differences in the activity of the microbiota might be responsible for the lack of response in those six patients. We therefore analyzed their metatranscriptomes at the time of diagnosis of BV (visit 1) and after treatment with metronidazole (visit 2) and compared them to those of 8 patients that responded to treatment according to Nugent score.

The high rate of recurrence is another crucial problem for BV treatment. The one-year recurrence rate of BV ranges from 40% to 80% after therapy with metronidazole (23) or clindamycin cream (24). CST dominated by *L. iners* might have an increased probability to shift to a dysbiotic state (22,25,26). In the second part of our study, we therefore followed the activity of the microbiota from four of the patients that initially responded to metronidazole treatment over a period of 3 months (visit 3–5) and analyzed gene expression of *L. crispatus* and *L. iners* in *vivo*.

We show the importance of *G. vaginalis* for BV, which can be massively underestimated using 16S rRNA gene sequencing. The relative abundance of the four subgroups of *G. vaginalis* could be determined in responders and non-responders. Transcripts potentially leading to lack of response to metronidazole treatment were identified. CRISPR-Cas genes are suggested as a novel mechanism of *G. vaginalis* to mitigate the DNA damaging effect of metronidazole. *L. iners* highly expressed genes for pore-forming toxins *in vivo*, and in *L. crispatus* the most highly expressed transcripts *in vivo* encoded enzymes for D-lactate and hydrogen peroxide production.

## Results

### Study population and overview of sequencing results

We studied the vaginal microbiome of 14 patients during and after metronidazole treatment of BV using metatranscriptome sequencing (Fig. 1A). Patients were part of a clinical trial described elsewhere (5). In the first part of the study, we analyzed samples from two timepoints (diagnosis of BV, visit 1) and after metronidazole treatment (visit 2). Eight patients responded to treatment, and six patients did not respond to treatment with the antibiotic, so were still BV positive according to Nugent score at visit 2. In the second part of our study, three additional timepoints were analyzed for four of the patients that initially responded to metronidazole therapy, covering a total period of 3 months. Those four patients belonged to the lactic acid arm of the clinical study. Two of them experienced recurrence, while the other two were stably non-BV after treatment according to Nugent score (details in Table S1 sheet 1). In total, we analyzed 40 vaginal fluid samples, 22 with BV status and 18 without. Metatranscriptome sequencing resulted in a total of 1,879,945,342 reads. Of these, 1,377,516,082 reads (73%) were left after quality filtering and removal of ribosomal RNA (Table S1 sheet 2). On average, 34 million reads were analyzed per sample.

**Fig. 1:**
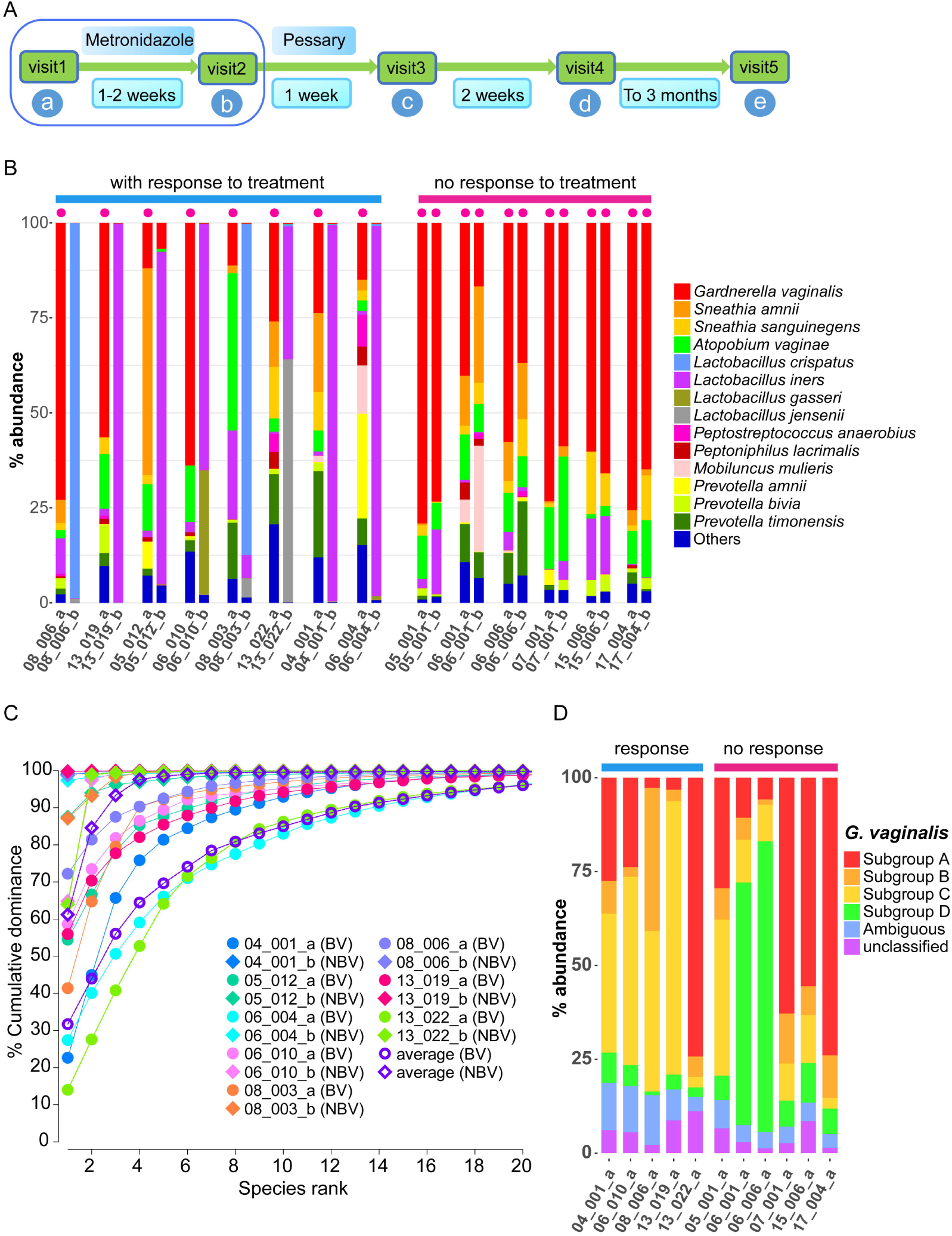
Study design and taxonomic composition of vaginal fluid metatranscriptomes in BV and after treatment with metronidazole. (A). Time course of the clinical study. (B) Taxonomic composition of the metatranscriptome at visit 1 (diagnosis) and visit 2 (after metronidazole therapy). (C) Cumulative dominance of the vaginal microbiota in BV and non-BV. (D) The subspecies composition of the *G. vaginalis* sub-community. Species with average relative abundance smaller than 0.5% were grouped into “Others”. The red dot on top of the samples indicates BV. The digits indicate the patient ID, while the letters a-b denote visit 1–2. After the first sampling at visit 1 the patients were treated with metronidazole. Total putative bacterial mRNA reads were mapped to the ref_Genome database using Kraken (see Methods for details). BV status was determined by Nugent score. In C, the “BV” and “NBV” in the parenthesis indicate BV and non-BV, respectively.

### Construction of the reference genome and gene databases for taxonomic and activity profiling

Human reads comprised ~11% (BV) versus ~56% (non-BV) of the total putative mRNA reads based on the standard Kraken (27) database (Table S1 sheet 2). This suggests that the bacterial load is much lower in non-BV than in BV since the human contamination is much higher in non-BV. Using the standard Kraken database, only 41% of total putative microbial (non-human) mRNA reads could be assigned taxonomically (Table S1 sheet 2). To improve the fraction of taxonomically assignable reads we then constructed a refined database (ref_Genome) which combined the urogenital subset of the HMP (28) database (147 genomes) and all species which are not included in the urogenital subset of the HMP database but detected by the standard Kraken database with an abundance >1% (7 genomes). We also added *S. amnii* and *S. sanguinegens* which had previously been shown to be highly abundant based on 16S rRNA gene sequencing (5) but were not contained in either HMP or the standard Kraken reference database. There are four *G. vaginalis* strains in the HMP database, of which one belongs to subgroup A and three belong to subgroup C. Given the importance and high intra-species diversity of *G. vaginalis*, we added 5 additional *G. vaginalis* genomes based on the genome tree reported in the NCBI database and the completeness of the genome assembly; these five strains cover all four subgroups. We added the genomes of *Gardnerella* sp. 26–12 and *Gardnerella* sp. 30–4 which were isolated from the bladder recently (29). They were classified into *G. vaginalis* subgroup A based on sequence homology (29). In total, this database contained 163 bacterial genomes from 105 species (Table S1 sheet 3). Using this database, the rate of taxonomically classified putative microbial mRNA reads could be improved to 86% on average (Table S1 sheet 2).

For functional assignment, we constructed a reference gene database (ref_Gene) (Table S1 sheet 4). It was based on the same genomes as the ref_Genome database, except that the seven additional *Gardnerella* spp. genomes were not included because of the low quality of the annotation of coding sequences. The ref_Gene database contained 301,323 genes. To investigate the activity shifts of the communities, we mapped the cleaned metatranscriptomic reads to the ref_Gene database using BWA. In total, 78% of total putative microbial mRNA reads could be mapped to the ref_Gene. Per sample, on average 8.9 million microbial mRNA reads could be mapped with MAPQ >10 (Table S1 sheet 2).

### Shifts in the taxonomic composition of the active community following metronidazole treatment

The taxonomic composition of transcripts was determined using Kraken and the ref_Genome database. Fig. 1B shows that in all communities with BV status the most abundant species were *G. vaginalis*, *A. vaginae*, *S. amnii* and *Prevotella timonensis* In the post treatment communities from responders (non-BV, Nugent score <6) the metatranscriptomes were dominated by *L. crispatus*, *L. iners* and *L. jensenii*, representing typical CSTs of the healthy female microbiota.

On average less than 14 species contributed >90% of the mapped reads in BV and 3 species accounted for >90% of the mapped reads in non-BV (Fig. 1C). The individual dominance plots showed the same pattern where 10 species contributed >90% of the metatranscriptomes for most patients in BV. In non-BV, this number was 2 for most patients and the dominance curves were extremely steep. For comparison, in the periodontal metatranscriptome more than 100 species were required to cover 90% of mapped reads (30). These data show that the active microbiota in BV is much less diverse than suggested by 16S rRNA gene sequencing.

*G. vaginalis* was the most dominant active species in BV. To estimate the relative abundance of the four subgroups of *G. vaginalis*, we extracted all reads assigned to *G. vaginalis* from the metatranscriptomes and assigned them to the four strains representing subgroups A-D, respectively (409–05, 00703Bmah, HMP9231, 00703Bmash) using Kraken. For this analysis, we used samples from visit 1 where *G. vaginalis* reads comprised at least 20% of all reads, which included all 6 patients without response to treatment, and 5 of the 8 patients that responded to treatment. Figure 1D shows that on average >95% of the reads could be mapped to the four subgroups and only on average 7% were assigned ambiguously. In those patients that did not respond to treatment, subgroups A and D comprised 68.5±17.2 % of all reads, while they accounted for 30.5±29.3% of all reads in patients that responded to treatment (Wilcoxon test P=0.0520). We observed that *Gardnerella* spp. previously isolated from the bladder (*Gardnerella* sp. 26–12 and *Gardnerella* sp. 30–4) (29) contributed on average 6% of all taxonomically assigned reads in BV (Fig. S1).

### Comparison of the taxonomic composition of vaginal fluid samples between metatranscriptome and 16S rRNA gene sequencing

We compared the taxonomic composition determined using 16S rRNA sequencing previously (5) and the taxonomic composition of the metatranscriptome determined here by Kraken with the ref_Genome database. In non-BV, we did not observe any considerable difference for the four most abundant species (Table S1 sheet 6), while in BV large differences between the two datasets were found. Fig. 2 shows the top 12 most abundant taxa identified using 16S rRNA gene sequencing and metatranscriptomics, respectively. Most of the abundant species identified in the mRNA sequencing data set were also identified using 16S rRNA gene amplicon sequencing, although usually at different abundances. For example, *A. vaginae* comprised 13% of all reads based on 16S rRNA gene sequencing, but only 11% in the metatranscriptome dataset. The most pronounced difference was observed for *G. vaginalis* which comprised on average 47% of relative abundance in the metatranscriptome and on average only 5% in the 16S rRNA sequencing data. Several additional differences were found: Higher level taxa like Veillonellaceae (family) or *Parvimonas* (genus) are not listed among the top 12 taxa of the metatranscriptome, because mapping occurred to species level, and the abundance of individual species of these higher order taxa was too low to be found among the top 12 taxa (Table S1 sheet 6). BVAB2 is readily detected by PCR and is an important indicator of BV, but it is not yet cultivated and so there is no genome available to map the reads against.

**Fig. 2:**
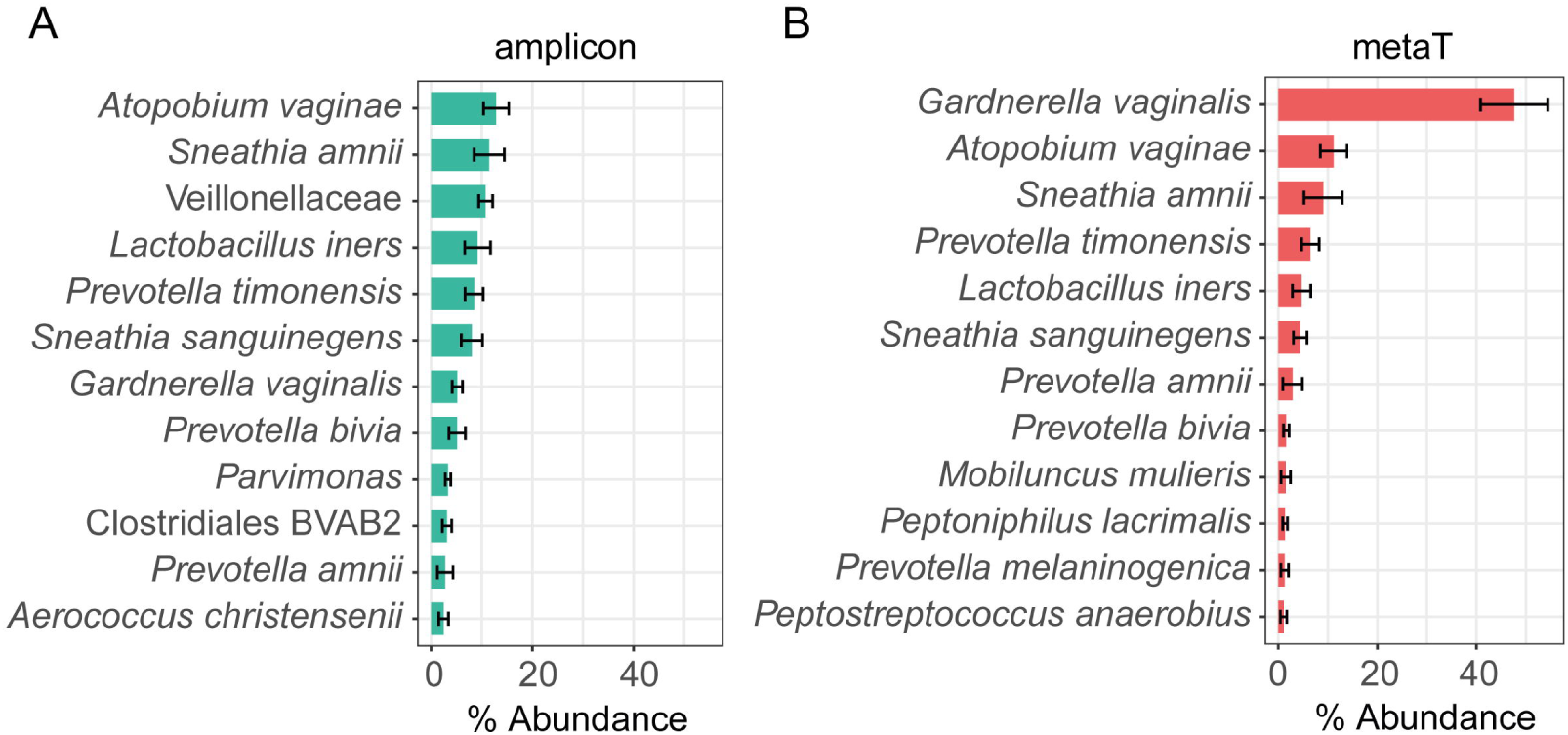
Average taxonomic composition of vaginal fluid samples in BV determined by 16S rRNA amplicon sequencing (A) and metatranscriptome sequencing (B). (A) Amplicon sequencing was performed as described in our previous study (5) using primers V1-V2. (B) The taxonomy was assigned based on all cleaned reads after removal of human reads using Kraken and the ref_Genome database. The top 12 most abundant taxa for each approach are shown in A and B. Relative average abundance was calculated based on all mapped reads. Mean and standard error are shown.

### Global community profiling in non-BV and BV

In order to profile the function of the communities, all cleaned putative mRNA reads (Table S1 sheet 2) were mapped using BWA onto the ref_Gene database annotated with KEGG ortholog (KO) genes. We used principal component analysis (PCA) to visualize the difference between the microbiota in BV and non-BV on the level of taxonomy (16S rRNA gene) (Fig. 3A), taxonomic composition of expressed genes (metatranscriptome) (Fig. 3B) and functional annotation of transcripts to KEGG orthologues (KO genes) (Fig. 3C). Figure 3A shows that the non-BV communities form a tight cluster on the level of the 16S rRNA sequencing, while BV communities vary, in accordance with the studies using amplicon sequencing of BV. *L. iners*, *Prevotella* spp., *G. vaginalis*, *A. vaginae* and *S. amnii* drive the separation between non-BV and BV On the level of the taxonomic composition of the metatranscriptomes (Fig. 3B) this pattern was reversed; samples from non-BV were much more heterogeneous than those from BV. The non-BV communities clustered into two groups dominated by *L. iners* and *L. crispatus* respectively, whereas *G. vaginalis*, *A. vaginae* and *S. amnii* were abundant in BV. This reversal is even stronger on the level of KO genes (Fig. 3C): Samples from BV form a tight cluster, while those from non-BV vary widely. It is the opposite pattern than that found for the phylogenetic marker gene. The KO genes that contributed most to these differences in non-BV were phosphofructokinase isozyme *pfkA* (31) and ribosomal protein coding genes *rpsI*, *rpmF* and *rplU* (32,33). In BV, *msmE*, *cycB* and *pflD* genes that encode proteins involved in carbohydrate uptake and metabolism (34,35) were stably higher expressed.

**Fig. 3:**
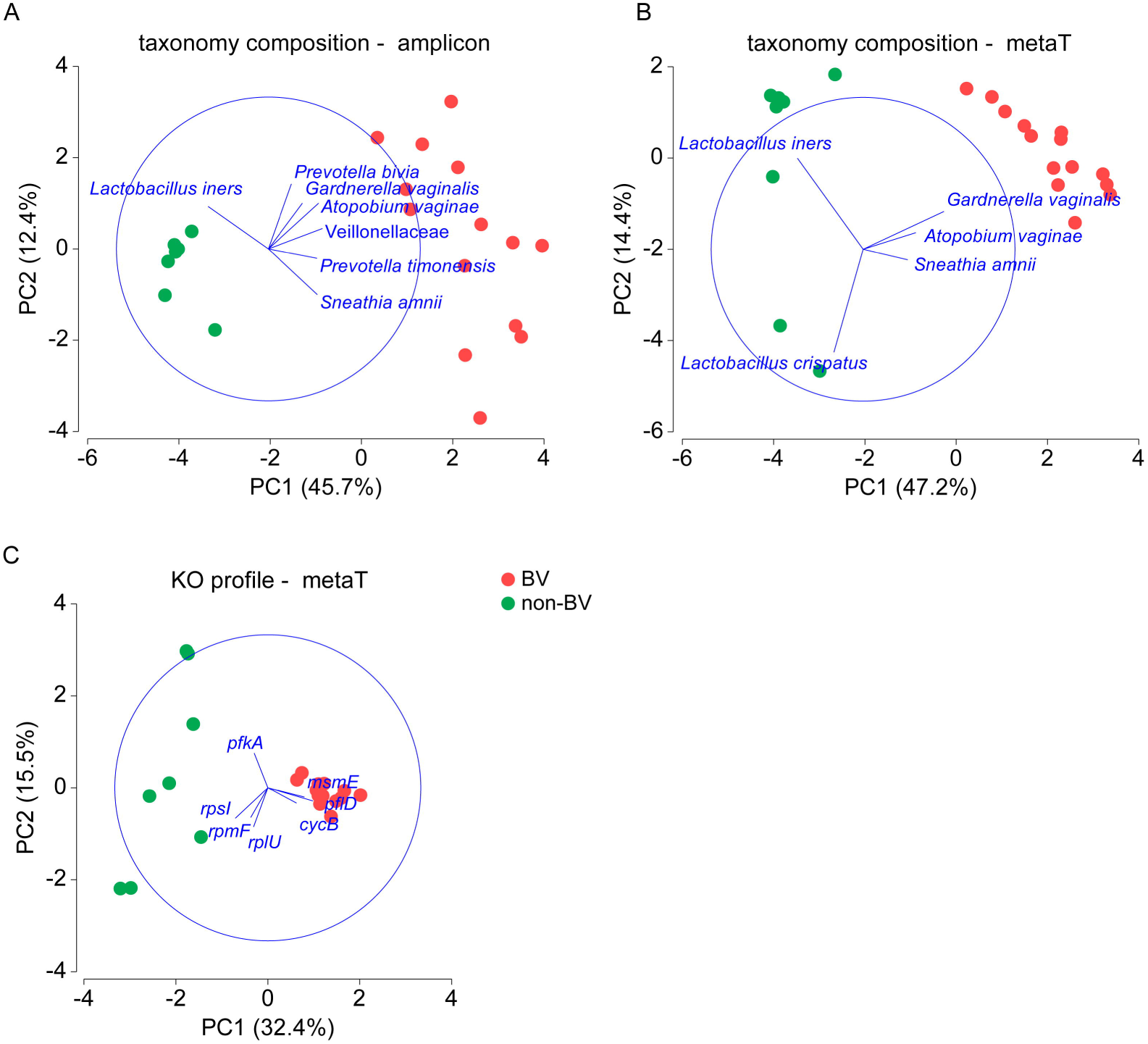
Principal components analysis (PCA) based on taxonomic profiles and activity profiles in BV and after metronidazole therapy (non-BV). (A) The PCA plot based on taxonomic profile using 16S rRNA gene sequencing. (B) The PCA based on taxonomic composition determined by metatranscriptome. (C) The PCA plot based on KO gene expression profile. The communities at visit 1 (BV) and visit 2 (after metronidazole therapy, non-BV) from 14 patients are shown. In the PCA biplots of taxonomy composition (Fig. 3A-B), the taxa with multivariate (multiple) correlation higher than 0.3 are illustrated, while for PCA (Fig. 3C) of KO profiles the KO genes with correlation >0.2 are shown.

### *In vivo* expression of putative metronidazole resistance associated genes in *G. vaginalis*

To clarify the possible contribution of genes related to metronidazole resistance in Gram positive pathogens to the difference in response to treatment of the vaginal microbiota, we examined their expression (Table S1 sheet 9) in *G*. *vaginalis*. For this analysis, BV communities from 11 patients at visit 1 were analyzed in which the level of *G. vaginalis* transcripts was >20%. Six of these patients did not respond to treatment and five responded. Although *A. vaginae* and *S. amnii* are also key players in BV we could not analyze them here since there were too few samples dominated by them. As shown in Fig. 4A, there was no clear expression pattern for most of these genes (detailed data in Table S1 sheet 9). The only significantly changed gene expression was that of the gene encoding ferredoxin which was less active in *G. vaginalis* in non-responding patients (fold change=1.67, Wilcoxon test P=0.00866 based on relative abundance = read count of given genes of *G. vaginalis* / read count of *G. vaginalis* %).

**Fig. 4:**
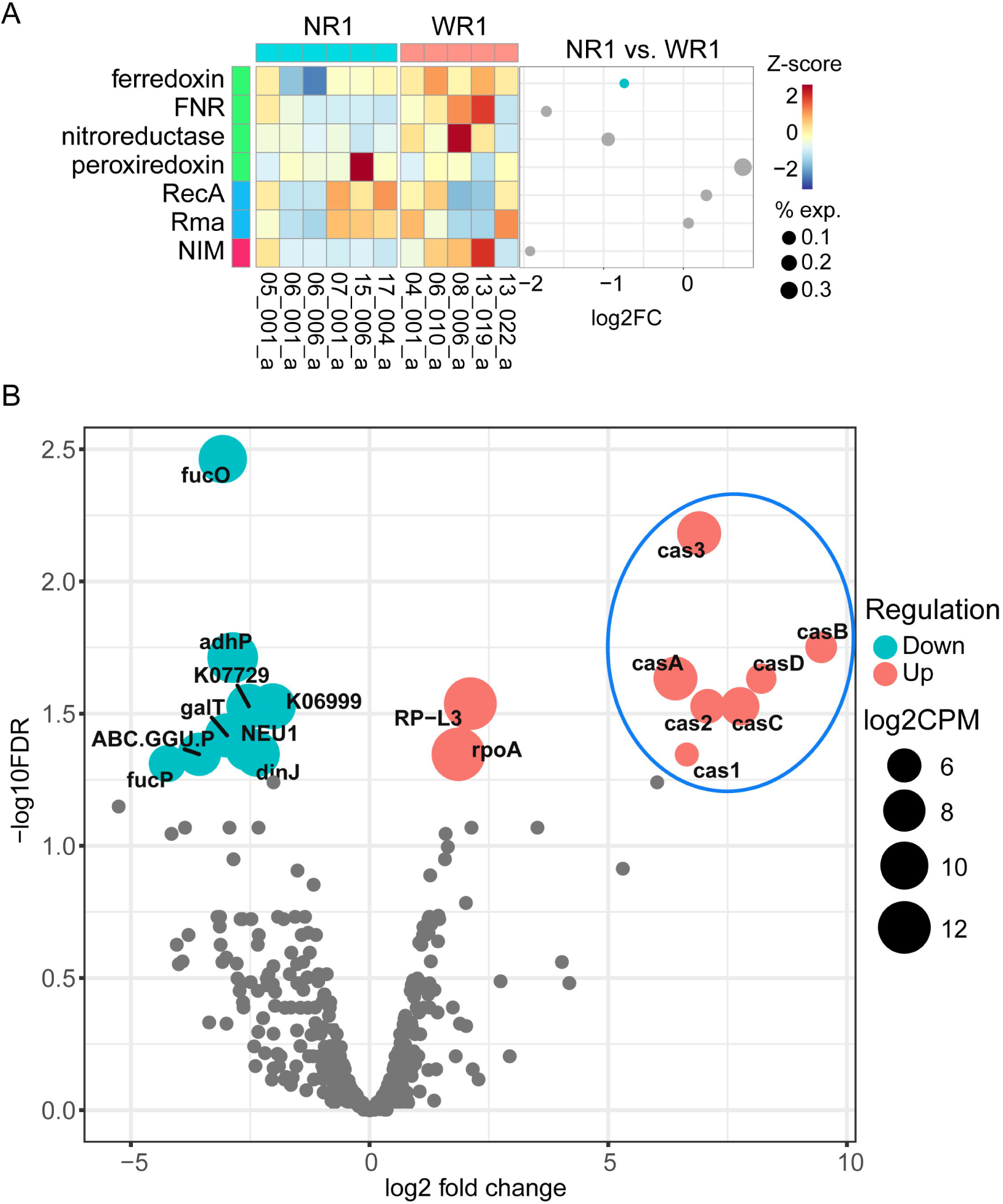
Changes in gene expression of *G. vaginalis* in patients responding to antibiotic treatment compared to non-responders. (A) Expression of putative metronidazole resistance associated genes of *G. vaginalis* in vaginal fluid microbiota. (B) Differential expression of KO genes: Seven *cas* genes of *G. vaginalis* were highly up-regulated in communities from patients who did not respond to the treatment. (A) The expression value was calculated based on relative abundance of reads mapped onto *G. vaginalis* using BWA. “NR1” (No Response 1) indicates the BV samples from six patients that did not respond to metronidazole treatment; “WR1” (With Response 1) represents the BV samples from four patients which afterwards responded to metronidazole. The dotplot illustrates the log2FC of the corresponding activity between *G. vaginalis* from non-responders and responders. The values in the heatmap were scaled using Z-sore. In the figure legend, “exp.” indicates the relative expression level. (B) “NR1” samples were compared with “WR1”. KO genes with FDR <= 0.05 are colored in red or turquoise (significantly differentially regulated) while FDR > 0.05 are in grey.

### CRISPR associated protein coding genes of *G. vaginalis* were strongly up-regulated in vaginal fluids of patients not responding to treatment

We then performed a global analysis of differential expression of KO genes of *G. vaginalis* in these same communities (visit 1, 11 BV samples with >20% transcripts from *G. vaginalis* including 6 patients that did not respond to treatment and 5 that responded). We observed that there were 9 KO genes highly up-regulated with FDR <= 0.05 (log2 fold change up to 9.46) in communities without response. Strikingly, among the most strongly up-regulated KO genes, seven were *cas* genes (36) (Fig. 4B). In total there were 8 different *G. vaginalis* CRISPR-associated (Cas) genes found in the genomes, namely *cas*1–3, *cas*A-E, of which seven were up-regulated (*cas*1–3, *cas*A-D).

There were 9 KO genes down-regulated but the fold change values were not as high as for the up-regulated genes. *fucP* (fucose permease) was identified as the most strongly down-regulated gene with a log2 fold change of −4.24.

### Time course of activity profiles and recurrence

In the second part of our study, we analyzed the metatranscriptome of vaginal fluid samples from four of the patients that initially responded to therapy with metronidazole for the complete duration of the clinical trial. Two of these patients experience recurrence of BV, and two remained stably non-BV. Five timepoints were analyzed of which the first two were already shown in Fig. 1A. They represented acute BV (visit 1) and after metronidazole therapy (visit 2). Here, we also show visit 3–5, which were all non-BV, with the exception of recurrence at visit 5 in patient 04_001 and at visit 3 in patient 06_004. Fig. 5A shows the taxonomic composition of the communities. In one of the two patients that stably maintained a non-BV status *L. crispatus*, and in the other *L. iners* dominated the microbiota. The principal components analysis of the activity profiles is shown in Fig. 5B. In acute BV, samples from all four patients clustered together (red circle). After the treatment, samples from patient 08_006 who was stably non-BV moved into a very dense and distinct cluster (illustrated by the arrow 1 and enclosed by a green circle). Samples from patient 13_019 who also remained stably non-BV moved to a different cluster after treatment, shown by arrow 2 and encircled blue.

**Fig. 5:**
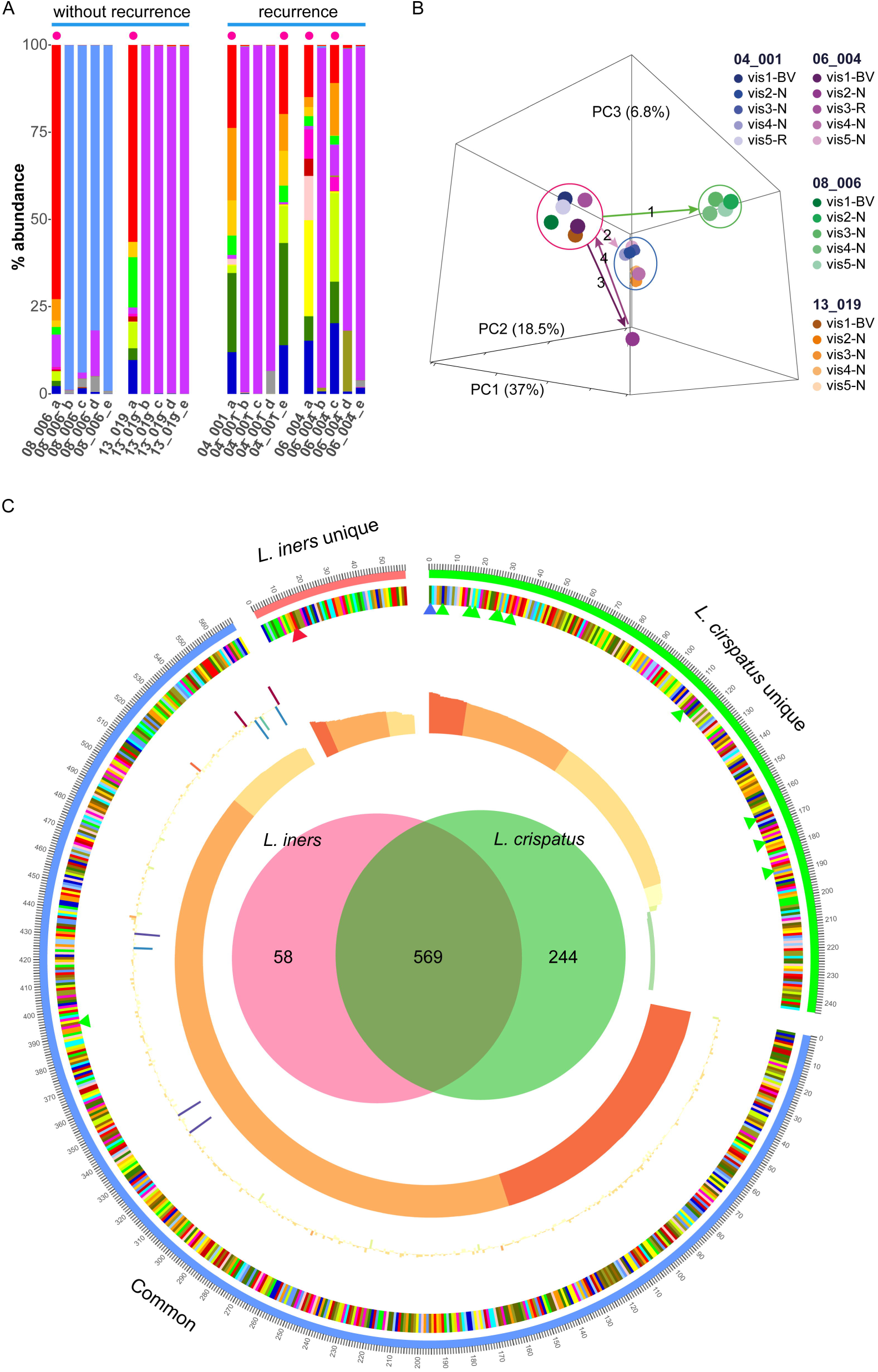
Shifts in the vaginal microbiome over 3 months. (A) Taxonomic composition of the metatranscriptome in two patients that were stably non-BV (without recurrence) and two patients that experienced recurrence. (B) PCA of activity profiles based on KO genes from the same patients. (C) Gene expression *in vivo* of *L. crispatus* and *L. iners*. (A) Acute BV and recurrence according to Nugent score are indicated as red dot. (B) Two women with recurrence (pink and blue color range) and two women without recurrence (green and orange color range) are shown. In the figure legend, BV indicates timepoint with BV, “R” indicates the recurrence and “H” represents health. The green and blue circles highlight healthy clusters, respectively, while the red circle highlights samples from BV. The arrows denote the temporal shifts of the communities during the treatment. (C) The Venn diagram indicates the unique KO genes of *L. crispatus* and *L. iners* as well as their shared KO genes. The innermost ring denotes the expression of KO genes by log2CPM, the outer ring illustrates the fold change of the expression of KO genes between *L. crispatus* dominated communities and *L. iners* dominated communities by log2FC. The KO genes are in descending order based on log2CPM. The small red triangles mark the inerolysin and hemolysin C genes, while blue and green triangles mark the genes encoding proteins involved in the production of D-lactic acid and hydrogen peroxide, respectively.

The activity shifts in patient 06_004, who experienced recurrence at visit 3, were especially noteworthy: After treatment, the community moved towards an activity profile distinct from all others (arrow 3). The recurrence of BV caused the community to shift back to the BV cluster (red circle, arrow 4). At visit 4, the community moved to the non-BV cluster (blue circle) and the patient became non-BV according to the Nugent score. We speculate that there was an unknown intervention after visit 3 which changed the microbiome but this was not recorded. Interestingly, the other case of recurrence (patient 04_001) had a different progression. From visit 2 to 4, patient 04_001 was non-BV and these samples clustered together in the “non-BV” cluster indicated by the blue circle. At visit 5, however, patient 04_001 had recurrent BV and the community shifted back again to the BV cluster.

### Transcriptomics of *L. iners* and *L. crispatus in vivo*

The stable colonization of the vaginal fluid of two patients that responded to antibiotic therapy with either *L. iners* or *L. crispatus* allowed us to profile their gene expression *in vivo* to gain more understanding of their different roles in the vaginal microbiota. We extracted the reads mapped on *L. crispatus* in patient 08_006 (timepoint b-e) and *L. iners* in patient 13_019 (timepoint b-e) and performed a differential expression (DE) analysis using edgeR to compare their activity profiles based on their KO genes comparing the expression of KO genes of *L. crispatus* in *L. crispatus* dominated samples (n = 4) with the expression of KO genes of *L. iners* in *L. iners* dominated samples (n = 4). The Venn diagram in Fig. 5C shows that the two species share 569 KO genes, while 58 are unique to *L. iners* and 244 are unique for *L. crispatus*, indicating *L. crispatus* possesses far more diverse functions than *L. iners*. The DE analysis identified 654 significantly differentially expressed KO genes, of which 393 were up-regulated in *L. crispatus* (Table S1 sheet 8). Among the top 100 most differentially expressed KO genes in terms of FDR value, 64 were up-regulated in *L. crispatus*, Remarkably, genes encoding enzymes involved in the production of H_2_O_2_ (pyruvate oxidase, NADH oxidase, glycolate oxidase) (37,38) were highly expressed in *L. crispatus* (Table S1 sheet 8, KO genes colored in blue). D-lactate dehydrogenase (K03778) was the most highly expressed gene in *L. crispatus* (log2 CPM = 13.3, Table S1 sheet 8, colored in light green) and this gene is absent in the genome of *L. iners*. On the other hand, we found that inerolysin (INY) was highly expressed (log2 CPM =9.8) in *L. iners*, but absent in the genome of *L*. *crispatus* (Table S1 sheet 8, colored in red). Interestingly, we also found the orthologous gene (K11031) of inerolysin known as vaginolysin (39) highly expressed in *G. vaginalis* (details in Table S1 sheet 7). Hemolysin C, another pore forming toxin, was highly expressed (log2 CPM = 9.8) in *L. iners* but absent in *L. crispatus*.

## Discussion

The aim of this study was to identify activity patterns in the vaginal fluid microbiota in BV and after metronidazole therapy. In particular, we compared the transcriptional profiles of *G. vaginalis* in vaginal microbiota from patients who did and did not respond to metronidazole treatment, respectively. This is the first study to investigate the activity alterations of the vaginal microbiota from patients with BV during treatment with the antibiotic metronidazole using the metatranscriptomics approach. We found several changes in gene expression in non-responding patients that might contribute to resistance against metronidazole by either not activating the pro-drug or repairing DNA damage.

*G. vaginalis* was the most dominant active species in BV. *G. vaginalis* can be divided into four phylogenetic subgroups which may in the future be described as subspecies and which differ in virulence and susceptibility to metronidazole (6,12). We found transcripts from all four subgroups in all patients, as previously shown based on sequencing of the universal target cp-60 gene (6,9,11). Interestingly, in those patients that did not respond to treatment, *Gardnerella* subgroups A and D which are resistant to metronidazole (14) were slightly more abundant.

Sequencing of phylogenetic marker genes like the 16S rRNA gene or the cpn60 universal target is a fast and sensitive method to profile the microbiota composition, but it does not provide functional information and is prone to PCR bias. Moreover, DNA from dead cells might also be detected. Therefore, we compared the taxonomic composition of the transcripts with that of the 16S rRNA genes determined previously in those samples (5). We observed that *G. vaginalis* comprised on average 47% of all transcripts in BV, while only 5% of 16S rRNA genes were assigned to this species. This suggests that *G. vaginalis* is transcriptionally more active than other vaginal bacteria; moreover, the commonly used 27F primer was previously shown to underrepresent *G. vaginalis* (40). Other differences between the two methods are caused by the low taxonomic resolution of the 16S rRNA gene, especially of short amplicons, where a large fraction of 16S rRNA reads is assigned to higher level taxa, e.g. genus or family. By contrast, the metatranscriptome reads are mapped to genomes and so have species level resolution. Finally, transcripts can only be mapped if a genome is available. If the species in question has not yet been cultivated, as for example the BVAB strains, then reads cannot be assigned. In the periodontal pocket microbiota, about 50% of all reads cannot be mapped to any bacterial genome (30). By contrast, the vaginal microbiota is much less diverse and most of its representatives have been cultivated; using the improved ref_Genome database, we were able to map 86% of all reads, indicating that uncultivated taxa did not contribute very significantly to the active community in BV and after metronidazole therapy.

Low diversity in health and high diversity in BV is a hallmark of BV and it is so striking that it has even been suggested to use diversity indices based on PCR amplified 16S rRNA genes in addition to the clinical diagnosis based on Amsel criteria and Nugent score (4,41–43). Our comparison between communities in BV and after metronidazole therapy on the levels of (1) 16S rRNA gene, (2) taxonomic composition of total transcripts, and (3) functional profiling based on KO genes shows a reversal of this observation: BV communities, although highly diverse on the taxonomic level, cluster tightly together on the functional level of KO genes. On the contrary, non-BV communities are similar on the taxonomic level, but highly diverse among individuals on the functional level.

In our metatranscriptome analysis we found evidence for mechanisms that hinder the activation of the metronidazole prodrug, or mitigate the damage that metronidazole inflicts on DNA, and thus could be important reasons for the lack of response in some women.

We show that the ferredoxin gene of *G. vaginalis* was less active in those patients that did not respond to metronidazole. As an electron carrier, ferredoxin is downregulated in *H. pylori* bacteria grown in the presence of metronidazole (17). It is required for activation of the prodrug in *H. pylori* (44) and might have a similar role in *G. vaginalis*. Lack of response might result from lack of activation of the prodrug. Unexpectedly, the nitroimidazole resistance (*nim*) gene, which has been shown to mediate resistance to metronidazole in *B. fragilis* by transforming metronidazole to a non-toxic amino derivative (16) was not highly expressed in non-responders. This could be due to technical problems, since the Nim protein sequence contains only partial CDS (https://www.ebi.ac.uk/ena/data/view/AGN03877). Moreover, *nim*-negative strains of *B. fragilis* can tolerate high levels of metronidazole indicating the importance of other mechanisms of resistance (16).

Remarkably, *cas* genes of *G. vaginalis* were highly up-regulated in samples from patients that did not respond to metronidazole treatment. The CRISPR-Cas genes are present in about half of all Bacteria and most Archaea (45); they represent a mechanism of adaptive immunity which protects the prokaryotic cell against foreign DNA and has been developed into a universal tool for genome editing (46). The *cas* genes of *G. vaginalis* belong to the *E. coli* subtype and were found in about half of the clinical isolates (36). Their up-regulation might reflect increased phage attacks in BV. Phages have been hypothesized to be crucial for the etiology of BV by causing the collapse of *Lactobacillus* populations (47); accordingly, *L. iners* upregulates its CRISPR-Cas system in BV (22). More than 400 annotated prophage sequences were found in 39 *Gardenerella* strains (29). They might be induced to enter the lytic cycle by the change in pH accompanying the shift to BV. However, the viral transcripts contributed 0.1% of the total metatranscriptome in both non-responders and responders before treatment.

The upregulation of CRISPR-Cas system genes in *G. vaginalis* from those patients that did not respond to treatment by metronidazole suggests that the CRISPR-Cas system might have a role in mitigating the DNA damaging effect of metronidazole. In addition to providing adaptive immunity, CRISPR-Cas systems can have various additional functions (48) and it was shown that they can protect the cell against DNA damaging agents (49). The Cas1 enzyme of *E. coli* (YgbT) physically and genetically interacts with the DNA repair system (RecBC, RuvB) and is recruited to DNA double strand breaks; moreover, YgbT is necessary for resistance of *E. coli* to DNA damage caused by the genotoxic antibiotic mitomycin C or UV light (49). Our findings suggest that the CRISPR-Cas system may protect the vaginal microbiota against the DNA damaging effect of metronidazole. If experimentally confirmed, this finding might open a new path to fight bacterial resistance against DNA damaging agents. For example, it would be worth testing if the susceptibility to metronidazole can be modified in *Gardnerella* isolates and possibly other vaginal pathogens according to the expression level of *cas* genes. It is not known how up-regulation of *cas* genes is regulated in the vaginal microbiota. It might be a response to phage attack, thus by suppressing *cas* genes the susceptibility to phages might be increased simultaneously with the susceptibility to metronidazole. Using CRISPR engineered phages for therapy of dysbiotic communities has been considered as one of many options of new therapeutic strategies based on a deeper understanding of the human microbiome (50).

*L. iners* and *L. crispatus* dominate their respective CST in the healthy vaginal microbiota. There were many factors observed by laboratory or genomic studies (37,51,52) which suggest that more protection against dysbiosis is provided by *L. crispatus* rather than by *L. iners*. Here we analyzed which genes are actually highly expressed *in vivo*: We observed that genes encoding proteins for the production of H_2_O_2_ and D-lactic acid were highly expressed in *L. crispatus*. H_2_O_2_ inhibits BV associated bacteria, but it has been questioned if its level in the vaginal milieu is high enough, and it was suggested that lactic acid is more protective (53). In *L. iners*, the genes for the pore-forming toxins inerolysin and hemolysin C were highly active, supporting the hypothesis that *L. iners* may play an ambiguous role in the vaginal econiche and is associated with vaginal dysbiosis (26,54).

## Conclusions

This first study of the *in vivo* transcriptional activity of vaginal fluid microbiota during metronidazole treatment of BV focused on possible reasons for the lack of response to antibiotic therapy in some patients. Genes related to activation of the prodrug and repairing the DNA damage caused by metronidazole were shown to be differentially expressed in responders and non-responders. A completely new role for Cas proteins is hypothesized which warrants closer inspection and may help to develop more efficient novel therapies to improve the treatment of BV.

## Material and Methods

### Study design

Vaginal fluid samples of women analyzed here were a subset of the samples obtained during a randomized controlled clinical trial described previously (5). The trial protocol was approved by the local ethics committee (Ärztekammer Nordrhein - Medical Association North Rhine) and written consent was obtained from all participants. The clinical trial was conducted in accordance with the Declaration of Helsinki on Ethical Principles for Medical Research Involving Human Subjects. Principles and guidelines for good clinical practice were followed. The study was registered on ClinicalTrials.gov with the identifier NCT02687789. Briefly, women were included into the clinical trial if they were BV positive according to Amsel criteria and Nugent score and were biofilm positive on vaginal epithelial cells and positive for extracellular polysaccharides (EPS) in urine. For treatment of acute BV, they received 2 g of metronidazole orally and were afterwards treated with an intravaginal pessary for three weeks, twice a week. Samples were taken during acute BV (visit 1), after receiving metronidazole 7 to 28 days after visit 1 (visit 2), after pessary application one week after visit 2 (visit 3), after continued pessary application two weeks after visit 3 (visit 4) and during follow up three months after visit 4 (visit 5).

The aim of the clinical trial had been to compare the effectiveness of two different types of pessary. The results and the taxonomic composition of the vaginal microbial communities have been reported (5). For the metatranscriptome analysis reported here, we chose a subset of 14 patients from the clinical trial. These 14 patients consisted of two groups named “with response to treatment” (n = 8) and “no response to treatment” (n = 6) (Fig. 1). Among the eight patients who responded to treatment, six had no recurrence and two experienced recurrence during the three months follow-up. For the analysis of lack of response to metronidazole, samples from all 14 patients were analyzed at two timepoints, acute BV (visit 1) and 7 to 28 days after antibiotic treatment (visit 2) (28 samples in total). For the analysis of recurrence, samples from all 5 visits were analyzed for 4 patients (two without recurrence and two with) (20 samples in total). These four patients all received the commercially available lactic acid pessary after metronidazole therapy at visit 3 and 4. Nugent score >6 was used to determine BV status since it is considered the gold standard for BV diagnosis (55) (Table S1 sheet 1). The BV status at visit 5 was determined by Amsel criteria as there was no Nugent score available at that timepoint. The sample ID was obtained by concatenating the patient ID and letters “a” to “e” indicating visit 1 to 5.

### Sample collection and transport

Vaginal fluid was obtained by infusing 2 ml of saline solution into the vagina followed by rotation against the vaginal wall with a speculum and then collecting the vaginal fluid with a syringe. Approximately 700 µl were immediately transferred to a tube containing 2 ml RNAprotect (Qiagen, Germany). Tubes were immediately frozen at −20°C, transported at −20°C within a week and stored at −70°C.

### RNA extraction and mRNA enrichment

RNA was extracted from 1 ml vaginal fluid suspension using the MO BIO PowerMicrobiome™ RNA Isolation Kit (Qiagen, Germany) with pretreatment: Vaginal fluid was centrifuged at 13,000 rpm for 1 minute. The pellet was resuspended in MoBio lysis buffer and this suspension was added to the supplied bead tubes filled with 500 µl ice-cold Phenol:Chloroform:Isoamyl Alcohol solution (Carl Roth, Germany). The bead-suspension mix was shaken at 5 m/s for 1 minute in 3 intervals which were 2 minutes apart using the MO BIO PowerLyzer™ (Qiagen, Germany). After centrifugation for 1 minute at 13,000 rpm and 4°C the upper phase containing the RNA was further processed according to the manufacturer’s instructions including DNAse I treatment. RNA was eluted in 100µl nuclease free water and vacuum concentrated to 50 µl. The Ribo-Zero Gold rRNA Removal Kit (Epidemiology) by Illumina (USA) was then used for mRNA enrichment with ethanol precipitation according to the manufacturer’s instructions. Integrity of RNA was evaluated using a Bioanalyzer 2100 (Agilent, Germany).

### Library preparation, sequencing and preprocessing of sequencing data

Paired-end mRNA Illumina sequencing libraries were constructed with the Script Seq Illumina Kit. Strand specific paired end sequencing was performed on the HiSeq 2500 Sequencer to yield 2 × 110 bp paired-end reads. Primers and sequencing adaptors were removed from raw sequencing data, followed by clipping the bases with quality score < 20 from the reads using Fastq-Mcf (56). After clipping, the remaining reads shorter than 50 were removed. Thereafter, the ribosomal RNA reads were eliminated using SortMeRNA v2.0 (57) with the default parameters.

### Taxonomy assignment using Kraken

Kraken (27), an accurate and ultra-fast taxonomy assignment tool for metagenomes was used to determine the taxonomic composition of the metatranscriptome data. Kraken uses the K-mer strategy and the lowest common ancestor (LCA) algorithm to affiliate a given read to a taxon. The standard Kraken database with addition of the human genome was used to identify human reads. This standard database consists of prokaryote genomes (2786), virus genomes (4418). The human genome (ver. GRCh38) was additionally downloaded from NCBI.

The ref_Genome database contained 163 bacterial genomes from 105 species of bacteria, including 147 genomes from the urogenital subset of the HMP reference genome sequence data (HMRGD). The complete list of reference genomes in the ref_Genome database can be found in Table S1 sheet 3. All results on the taxonomic composition in this study were achieved based on this database.

### Short reads alignment by BWA

Kraken was used for taxonomy classification, while BWA was applied to determine the expression of genes. A reference gene database named ref_Gene was constructed which contained the genes from the urogenital tract subset of the HMRGD and the genes from 9 additional genomes (Table S1 sheet 4). The genes of *Gardnerella sp*. 26–12, *Gardnerella sp*. 30–4 could not be included because only less than half of their CDS are available (around 1000 genes). The short reads alignment was performed using BWA with the BWA-MEM (58) algorithm. A mapping seed length of 31 which is much longer than the default seed length 19 was applied to achieve reliable alignments. Reads that mapped with mapping quality score (MAPQ) lower than 10 were excluded. MAPQ contains the Phred-scaled posterior probability that the mapping position is wrong (59).

### KEGG Orthologous (KO) gene annotation of ref_Gene database

The ref_Gene database was annotated using KEGG prokaryote protein sequences. The KEGG prokaryote protein sequence database represents a non-redundant protein dataset of Bacteria and Archaea on the species level and contains about 7 million non-redundant peptide sequences grouped into 14,390 distinct KO genes. A KO gene contains several genes from different species with similar function. DIAMOND (60), a much faster alternative to BLASTX was applied to map the ref_Gene sequences against the KEGG prokaryote protein sequence database with its “more sensitive mode”. To obtain reliable annotation, only alignments with sequence identity >=50 and E-value <= 1e-5 and query coverage >= 70% were taken into account. By annotating the genes in the ref_Gene database to KO genes, we were able to determine the expression profile of KO genes, and investigate the activity shifts in BV based on differential expression analysis of KO genes. The cumulative dominance analysis and PCA were carried out using Primer 7 (61).

### Differential expression (DE) analysis

All differential expression (DE) analyses were performed using the R package edgeR (62). The Benjamini Hochberg (BH) method was used to correct the p value of DE analysis with the false discovery rate (FDR) for multiple comparisons. Genes with FDR smaller than 0.05 were considered as significantly differentially regulated. The sample groups defined for each comparison are listed in Table S1 sheet 1.

### Detection of putative metronidazole resistance related genes

To detect the expression of previously reported putative metronidazole resistance genes such as recA, recA-mediated autopeptidase (Rma), peroxiredoxin, nitroimidazole resistance protein (NIM), ferredoxin/ferredoxin-NADP reductase (FNR), nitroreductase and ferredoxin, we examined the expression level of these genes for *G. vaginalis* in the vaginal community from patients without response to metronidazole treatment (n = 6) as well as with response (n = 4). As most of these genes do not have corresponding KO genes, we annotated the ref_Gene database based on the sequences of these genes using BLASTN. The sequences were retrieved from ENA by key words of each “ferredoxin, NADPH flavin oxidoreductase, nitroreductase, peroxiredoxin, pyruvate ferredoxin oxidoreductase, recA, nitroimidazole resistance” plus *G. vaginalis*. In total, 155 unique sequences of *G. vaginalis* were obtained for the annotation of ref_Gene database. The identification of duplicate sequences was done by SeqKit (63).

### Availability of data and material

The sequencing data have been deposited in the European Nucleotide Archive with accession number PRJEB21446.

## Authors’ contributions

This study was designed by IWD and CG. CM and CA provided the clinical samples. RNA extraction and mRNA enrichment was performed by CG. SB prepared cDNA libraries, and performed Illumina sequencing. Z-LD performed all the data analyses. Data interpretation and visualization were done by IWD, Z-LD and CG. Z-LD and CG wrote the manuscript draft, and all authors reviewed the manuscript.

## Acknowledgements

This work was supported by ZIM project grant KF3134201MD3 of the Bundesministerium für Wirtschaft und Energie (BMWi), Germany. The clinical study was funded by Dr. August Wolff GmbH & Co. KG Arzneimittel within the ZIM project KF3134201MD3.

## Supplementary Information

**Fig. S1: Taxonomic composition of communities on the species level determined by metatranscriptome sequencing in non-BV and BV.** The species present in at least 2 samples with relative abundance >1% are shown. The *Gardnerella* bladder isolates are illustrated separately from *G. vaginalis*. The sample name in red indicates the first BV incidence of patients with recurrence and purple depicts the second incidence.

**Supplementary Table S1: All supplementary data**

**Sheet 1: Sample description**

**Sheet 2: Read summary**

**Sheet 3: Genomes in the ref_Genome database for taxonomic assignment**

**Sheet 4: Genomes in the ref_Gene database with a total of 301,323 genes for functional assignment with BWA**

**Sheet 5: Species composition determined by Kraken based on the ref_Genome database for taxonomic assignment**

**Sheet 6: Comparison of the community composition determined by 16S rRNA gene amplicon sequencing (V1-V2) and metatranscriptome sequencing**

**Sheet 7: Gene expression based on the ref_Gene database for functional assignment with KO annotation**

**Sheet 8: The differential expression of KO genes between *L. crispatus* and *L. iners***

**Sheet 9: Expression of metronidazole activation and resistance associated genes in *G. vaginalis***

**Sheet 10: Differential expression of KO genes of *G. vaginalis* from communities without response to metronidazole treatment compared to those with response**

